# Type I and III interferons synergize with TNF to promote virally-triggered damage to the intestinal epithelium

**DOI:** 10.64898/2026.01.10.698817

**Authors:** Lucie Bernard-Raichon, Jessica A Neil, Kayla Kim, Thomas Heaney, Brittany M Miller, Damee Moon, Ashira Lubkin, Ashley L DuMont, Victor J Torres, Jordan Axelrad, Yu Matsuzawa-Ishimoto, Ken Cadwell

**Affiliations:** Kimmel Center for Biology and Medicine at the Skirball Institute, New York University Grossman School of Medicine, New York, NY, USA; Department of Microbiology, New York University Grossman School of Medicine, New York, NY, USA; Division of Gastroenterology and Hepatology, Department of Medicine, University of Pennsylvania Perelman School of Medicine, Philadelphia, PA, USA; Institut de Génomique Fonctionnelle de Lyon, École Normale Supérieure de Lyon, CNRS UMR 5242, UCBL Lyon-1, Lyon F-69007, France; Department of Microbiology and Immunology, Peter Doherty Institute for Infection and Immunity, University of Melbourne, Melbourne, VIC, Australia; Institute for Systems Genetics, New York University Grossman School of Medicine, New York, New York, United States; Fungal Pathogenesis Section, Laboratory of Clinical Immunology and Microbiology, National Institute of Allergy and Infectious Diseases (NIAID), National Institutes of Health (NIH), Bethesda, MD, USA; Postdoctoral Research Associate program, National Institute of General Medical Sciences, National Institutes of Health (NIH), Bethesda, MD, USA; Department of Host-Microbe Interactions, St. Jude Children’s Research Hospital, Memphis, TN, USA; Division of Gastroenterology, Department of Medicine, NYU Grossman School of Medicine, New York, USA; Department of Clinical and Diagnostic Laboratory Science Graduate School of Medical and Dental Science Institute of Science Tokyo (Science Tokyo), Tokyo, Japan; Department of Pathobiology, University of Pennsylvania School of Veterinary Medicine, Philadelphia, PA, USA

**Keywords:** Norovirus, interferons, autophagy, intestinal inflammation, necroptosis, Paneth cells

## Abstract

Excessive cell death in the epithelium due to prolonged immune activation is associated with intestinal diseases such as Crohn’s disease. Mice with mutations in the Crohn’s disease susceptibility gene *Atg16l1* are susceptible to inflammation associated with intestinal epithelial cell (IEC) death, including loss of antimicrobial Paneth cells triggered by infection with murine norovirus (MNV). Here, we show that intestinal disease downstream of MNV depends on IFN-α/β and IFN-λ signaling in IECs. In mouse organoids, IFNs synergize with TNF to induce RIPK1-dependent cell death, amplified by ATG16L1 deficiency. We further show that human intestinal organoids harboring the *ATG16L1* risk allele exhibit heightened sensitivity to TNF and IFN co-stimulation and to serum from severe COVID-19 patients. Our findings reveal that virally triggered cytokines including IFNs exacerbate epithelial vulnerability in genetically predisposed hosts.

## Introduction

A single layer of epithelial cells in the intestine forms the first line of defense against diverse microbial agents ranging from beneficial members of the microbiota to pathogens that cause life-threatening disease^1^. Intestinal epithelial cells (IECs) include enterocytes that absorb nutrients, chemosensory tuft cells, mucus-secreting goblet cells, and enteroendocrine cells that produce hormones. The small intestinal crypt also harbors Paneth cells, IECs that secrete antimicrobial factors such as lysozyme and defensins. The proper turnover of IECs maintains anatomical integrity and contributes to resolving infections, such as through sloughing of damaged or infected cells^2^. However, inflammatory or excessive cell death triggered by prolonged immune activation is a hallmark of inflammatory bowel diseases (IBDs) such as Crohn’s disease (CD) and ulcerative colitis^3^. The cell death modality, timing, and cell type specificity are likely important variables that contribute to outcomes in infectious and inflammatory diseases of the gut.

The etiology of IBDs is complex and associated with genetic and environmental factors^4^. Although sustaining remission continues to be a challenge in many patients, the success of therapies that target cytokines and their signaling activities support a disease model in which these soluble immune mediators disrupt the epithelial barrier in a susceptible host. Consistent with this possibility, we demonstrated that mutations in the CD susceptibility gene *ATG16L1* are associated with loss of function and viability of Paneth cells as well as susceptibility to intestinal injury downstream of TNF^5–8^. ATG16L1 is necessary for the cell biological process of autophagy in which cytosolic material, such as protein aggregates and damaged organelles, is targeted to the lysosome for degradation and recycling^9^. ATG16L1 and other autophagy genes have an epithelial-intrinsic role in promoting intestinal barrier function^7, 10–16^. Autophagy and related processes may be particularly important in Paneth cells because of their longevity (2-3 months) and high secretory burden that make these cells susceptible to ER and mitochondrial stress^5, 7, 17, 18^.

We previously developed a model in which *Atg16l1*-mutant mice, including mice with an IEC-specific deletion of *Atg16l1* (*Atg16l1*^fl/fl^;villin-cre, hereafter referred to as *Atg16l1*^ΔIEC^), persistently infected with murine norovirus (MNV) are susceptible to chemical injury to the gut by dextran sodium sulfate (DSS) and develop Paneth cell alterations similar to those observed in patients with CD homozygous for the *ATG16L1 T300A* risk variant^6, 7^. Virally-triggered disease outcomes are dependent on TNF^6, 7^. This IEC-intrinsic role for ATG16L1 in mitigating damaged caused by TNF is reproduced in small intestinal organoids, a 3D cell culture model differentiated from intestinal stem cells. We and others showed that organoids generated from *Atg16l1*-mutant mice or patients with *ATG16L1^T300A^*undergo necroptosis (programmed necrosis) and loss of Paneth cells upon TNF treatment^7, 13^. Given that MNV can promote beneficial immune development under certain settings and is otherwise tolerated^19–21^, the virally-triggered intestinal damage observed in *Atg16l1*^ΔIEC^ mice provides an opportunity to identify the immune factors that regulate IEC-viability in response to environmental perturbations.

Type I Interferons (IFN-α/β) and type III interferons (IFN-λs) are best known for their antiviral functions downstream of Janus kinases (JAK) and signal transducer and activator of transcription proteins (STAT) upon receptor engagement. Although type I and III IFNs induce similar interferon stimulated genes (ISGs), they have distinct activities mediated in part by differential expression of their receptors. In the intestine, the type I IFN receptor (IFNAR) primarily functions in immune cells and is downregulated in IECs in adult mice whereas the type III IFN receptor (IFNLR) is highly expressed in IECs and neutrophils^22–24^. In the context of MNV infection, IFN-α/β prevents extraintestinal dissemination while IFN-λ restricts local replication and fecal shedding^25–29^. Although homeostatic IFN-λ produced by lymphocytes in response to commensal bacteria limits viral replication during early infection^30^, the viral non-structural protein NS1 encoded by persistent strains mediates evasion of IFN-λ to establish prolonged infection of tuft cells^31, 32^. In addition to their antiviral role, IFNs impact intestinal homeostasis. A recent study reported that mice lacking both IFNAR and IFNLR do not survive even low levels of DSS exposure, which was associated with extensive destruction of the colonic epithelium^33^. By contrast, IFN-λ levels are high in serum and small intestinal tissue of patients with CD, and over-expression or administration of IFN-λ induces inflammatory forms of epithelial cell death, necroptosis and pyroptosis^34, 35^.

Although MNV can activate cell death proteins directly in infected IECs to promote its reproduction and immune evasion^36–41^, persistent strains display specific tropism for tuft cells due to the restricted expression of the viral entry receptor *Cd300lf* ^31, 42, 43^. Thus, in the *Atg16l1* mutant setting, in which cell death is observed in Paneth cells that do not express *Cd300lf*, extracellular factors mediate IEC defects. MNV increases TNF and IFN-γ producing intra-epithelial lymphocytes and inhibits protective γδ T cells upstream of Paneth cell death and DSS-induced disease^6–8^. Here, we identify a critical cytotoxic role of type I and III IFNs in mediating these effects. We report that deletion of IFNAR or IFNLR in IECs rescues intestinal defects in *Atg16l1* deficient mice and identify a role for IFNs in potentiating TNF-induced cell death. Using organoids and serum from patients with COVID, we provide evidence that the general epithelial cytotoxic effect of cytokines produced in response to viral infections is conserved in humans. Thus, our results suggest that a component of viral sequelae includes production of soluble immune mediators such as TNF and IFNs that compromise the viability of IECs, which is especially detrimental in genetically susceptible hosts.

## Results

### Epithelial type I and III interferon signaling mediate intestinal damage downstream of viral infection

*Atg16l1*-mutant mice display spontaneous ISG expression in the gut dependent on the microbiota^44, 45^. To determine whether ISG expression is further increased in the presence of MNV, we examined the small intestine (ileum) and colon of WT and *Atg16l1^ΔIEC^* mice 10 days post intragastric infection with MNV strain CR6. The levels of three representative ISGs (*Mx2*, *Oasl2*, and *Zbp1*) in the ileum and colon were increased by MNV infection, and for the ileum, highest in MNV-infected *Atg16l1^ΔIEC^* mice (Sup. Fig. 1A). IFN-λ is primarily responsible for ISG expression in IECs^46^. To confirm that IFN-λ is responsible for the ISG expression, we crossed *Ifnlr* floxed mice to *Atg16l1^ΔIEC^* mice to obtain *Atg16l1;Ifnlr^ΔIEC^*mice that are deficient in ATG16L1 and IFNLR in IECs. ISG expression in MNV-infected *Atg16l1;Ifnlr^ΔIEC^*mice was not significantly different from uninfected Cre-negative controls, hereafter referred to as wild-type (WT) (Sup. Fig. 1A).

To determine whether IFN-λ signaling contributes to virally triggered intestinal defects, we examined the susceptibility of *Atg16l1;Ifnlr^ΔIEC^*mice to DSS following MNV infection. *Atg16l1^ΔIEC^* mice showed increased lethality, weight loss, and disease score compared to WT mice upon MNV + DSS treatment (Fig. 1A-C), similar to our previous study^7^. In contrast, these disease signs were abrogated in MNV + DSS treated *Atg16l1;Ifnlr ^ΔIEC^* mice and were similar to *Ifnlr^ΔIEC^* (*Ifnlr*^fl/fl^villin-cre) and WT controls (Fig. 1A-C). We also generated *Atg16l1;Ifnar^ΔIEC^* (*Atg16l1^fl/fl^Ifnar*^fl/fl^villin-cre) mice for additional comparison and unexpectedly found that disease was ameliorated. Consistent with a dominant antiviral role of IFNLR at the epithelial barrier during MNV infection^26^, *Atg16l1;Ifnlr^ΔIEC^* mice, but not *Atg16l1;Ifnar^ΔIEC^*mice, displayed increased shedding of virus in stool compared to *Atg16l1^ΔIEC^* or WT mice (Fig. 1D). None of the mice examined displayed mortality or exacerbated disease in response to DSS in the absence of MNV, except *Atg16l1;Ifnlr^ΔIEC^*mice (Sup. Fig 1B). These results indicate that IFNLR and IFNAR in IECs are necessary for the exacerbated intestinal injury response in *Atg16l1 ^ΔIEC^*mice infected with MNV and may have a different role in uninfected *Atg16l1* mutant mice.

**Figure 1:**
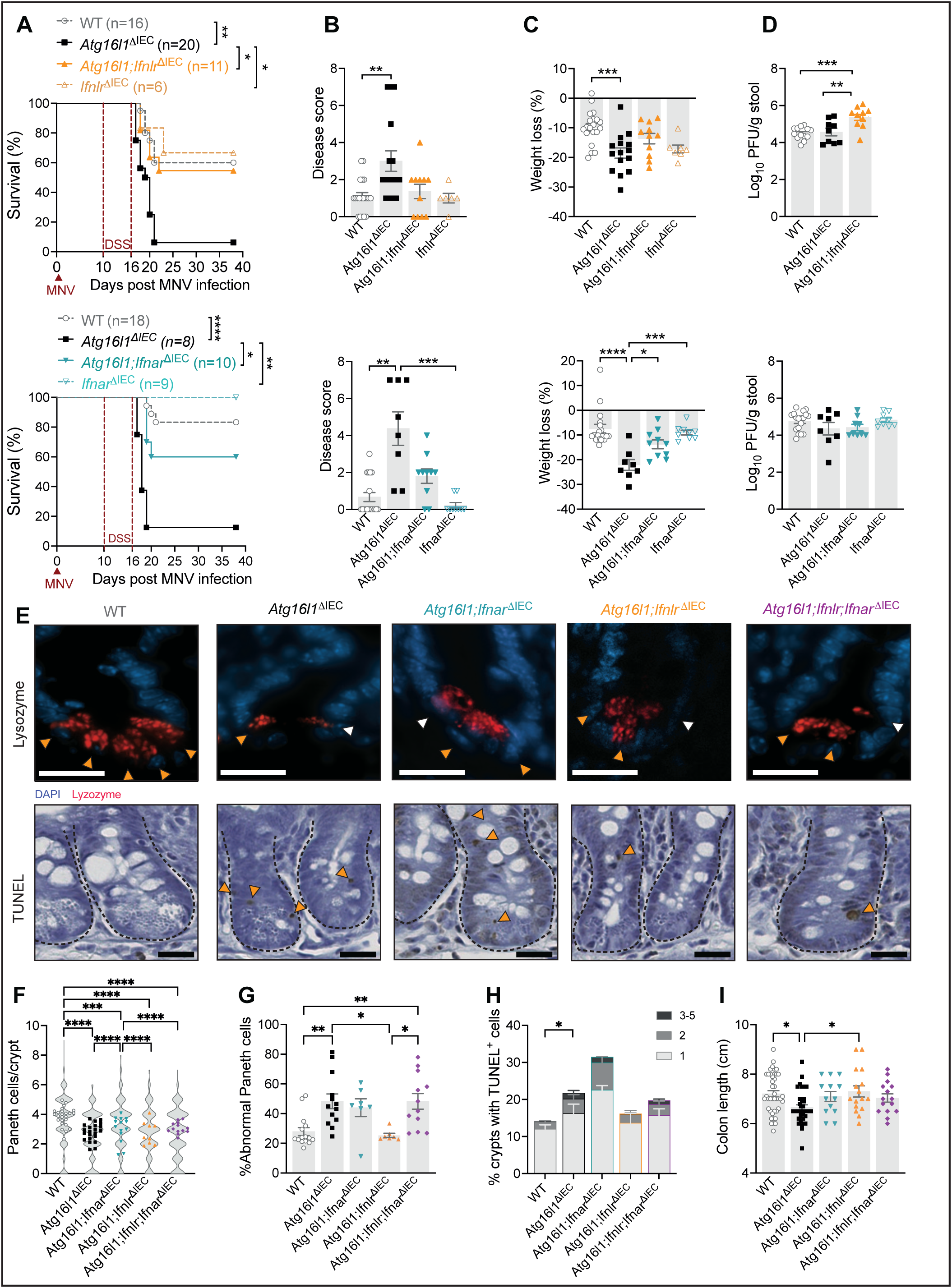
Epithelial type I and III interferon signaling mediate intestinal damage downstream of viral infection. **(A)** Survival of wildtype (WT; villin-cre^−^ littermate controls), *Atg16l1*^Δ*IEC*^, *Atg16l1;Ifnlr*^Δ*IEC*^, *Ifnlr*^Δ*IEC*^, *Atg16l1;Ifnar*^Δ*IEC*^, and *Ifnar*^Δ*IEC*^ mice receiving 5% DSS in drinking water for 6 days starting 10 days post inoculation (dpi) with MNV. **(B-C)** Cumulative disease score (B) and weight loss (C) at day 6 post DSS from mice in (A). **(D)** Plaque forming units (PFUs) of MNV per gram of stool from mice in (A) at 10 dpi. **(E-F)** Representative images of ileal crypts stained for lysozyme (red) and DAPI (blue) (top) or for deoxynucleotidyl transferase–mediated dUTP nick-end labeling (TUNEL) (bottom) of WT, *Atg16l1^ΔIEC^*, *Atg16l1;Ifnar*^Δ*IEC*^, *Atg16l1;Ifnlr*^Δ*IEC*^, and *Atg16l1;Ifnlr;Ifnar*^Δ*IEC*^ mice infected with MNV on day 5 post DSS treatment as in (A) (scale bars 20μm). **(F)** Violin plots depicting the repartition of the number of Paneth cells observed in individual crypts, overlayed with symbols representing the average number of Paneth cells per crypt observed in each mouse based on the lysozyme staining shown in (E). **(G)** Average proportion of abnormal Paneth cells per crypt observed in each mouse as determined on the basis of whether each Paneth cell displayed a typical staining pattern with distinguishable granules (normal, orange arrow) or depleted and/or diffuse lysozyme staining (abnormal, white arrow) as indicated in (E). **(H)** Proportion of crypts with 1, 2 or 3-5 TUNEL+ cells per mouse of each genotype. Orange arrowheads indicate TUNEL^+^ cells in (E). At least 50 villi and crypts were quantified per mouse. **(I)** Measurement of colon length from mice in (E-H). Each symbol represents individual mice (or the mean in A) with bars representing mean ± SEM from at least two independent experiments. Comparison between all genotypes was performed, not significant comparisons are not indicated, *p≤0.05, **p≤0.01, ***p≤0.001, ****p≤0.0001, See also Figure S1.

We previously showed that Paneth cells are less frequent and structurally altered in the ileum and that colons are shortened in MNV + DSS treated *Atg16l1^ΔIEC^* mice^7^. *Atg16l1^ΔIEC^* mice also displayed a lower number of Paneth cells compared to similarly-treated WT mice, and the remaining Paneth cells showed diffuse lysozyme staining and/or reduced number of lysozyme^+^ granules that are a characteristic of CD^6^. Deleting *Ifnlr* or *Ifnar* in *Atg16l1 ^ΔIEC^* mice led to partial reversal of Paneth cell abnormalities. The number of Paneth cells in *Atg16l1;Ifnar^ΔIEC^* mice were increased compared to *Atg16l1^ΔIEC^* mice and the proportion of abnormal Paneth cells were reduced in *Atg16l1;Ifnlr^ΔIEC^*mice (Fig. 1E-G). To test potential redundancy, we crossed *Atg16l1;Ifnlr^ΔIEC^* mice with *Atg16l1;Ifnar^ΔIEC^* mice to generate *Atg16l1;Ifnlr;Ifnar ^ΔIEC^* mice deficient in both receptors. However, *Atg16l1;Ifnlr;Ifnar ^ΔIEC^* mice still displayed reductions in Paneth cells and a high proportion showed abnormal lysozyme staining compared with WT mice (Fig. 1E-G). We cannot rule out the possibility that the above analyses would miss Paneth cells that lose their granules, which can happen in the *Atg16l1* mutant setting^5^. To more broadly capture abnormalities, we used terminal deoxynucleotidyl transferase–mediated dUTP nick-end labeling (TUNEL) to detect cell death. TUNEL-positive cells were rarely detected in the crypts of WT mice whereas a subset of crypts displayed multiple TUNEL-positive cells in *Atg16l1 ^ΔIEC^*mice (Fig. 1E, H). Rather than a decrease, the number of TUNEL-positive cells displayed a non-significant increase in *Atg16l1;Ifnar ^ΔIEC^* mice compared with *Atg16l1 ^ΔIEC^* mice. However, the *Atg16l1;Ifnlr ^ΔIEC^* mice were similar to WT controls and *Atg16l1;Ifnlr;Ifnar ^ΔIEC^* mice were similar to *Atg16l1 ^ΔIEC^* mice (Fig. 1E, H). *Atg16l1^ΔIEC^* mice also exhibited increased colon shortening compared to WT and *Atg16l1;Ifnlr ^ΔIEC^* mice following MNV + DSS (Fig. 1I).

These results indicate that IFNLR and IFNAR signaling mediate lethality in MNV-infected *Atg16l1^ΔIEC^* mice. IFNLR and IFNAR contribute to Paneth cell abnormalities and cell death, but deleting these receptors individually or together do not fully reverse these IEC defects.

### IFNβ and IFNλ exacerbate TNF-mediated cell death of intestinal organoids

We previously found that organoids generated from *Atg16l1^ΔIEC^*mice (*Atg16l1^−/−^* organoids) display a striking spontaneous ISG signature and that treatment with the JAK inhibitor ruxolitinib prevents TNF-induced cell death of these *Atg16l1^−/−^* organoids^10^. Although these results are consistent with a role for signaling downstream of cytokine receptors that use the JAK/STAT pathway, it was unclear whether IFNs can cooperate with TNF to induce loss of organoid viability. Since the majority of *Atg16l1^−/−^*organoids die within the first 2-3 days of seeding when TNF is administered on day 0^7^, we modified our protocol to increase the dynamic range of the assay and assess the synergistic effect of TNF and IFNs. We found that >50% of *Atg16l1^−/−^* organoids and ∼80% of *Atg16L1^+/+^* organoids (derived from *Atg16l1*^fl/fl^ mice that were negative for Cre) remained viable 72 hours post-stimulation when TNF was added 3 days after seeding (Fig. 2A, B). Treatment with IFN-β and IFN-λ2 led to similar or slightly greater loss of viability as TNF for both *Atg16L1^+/+^*and *Atg16l1^−/−^* organoids (Fig. 2A, B). In contrast, the combination of TNF and IFN led to substantial organoid death. After only 24 h, *Atg16l1^−/−^* organoids stimulated with TNF + IFN-β or TNF + IFN-λ2 showed complete loss of viability (Fig. 2A, B). *Atg16L1^+/+^* organoids started to display loss of viability at 24 h and were completely dead by 48-72 h post-treatment with TNF + IFN-β or TNF + IFN-λ2 (Fig. 2A, B). Titration of IFN-β and IFN-λ2 in the presence of TNF showed dose responsiveness and that *Atg16l1^−/−^* organoids are more sensitive to IFN since 10x less IFN induced same level of cell death in this genotype compared with *Atg16L1^+/+^* organoids (Fig. 2C). Therefore, IFN-β and IFN-λ2 increase the toxicity of TNF, and ATG16L1 deficiency further sensitizes organoids to the combination of these cytokines.

**Figure 2:**
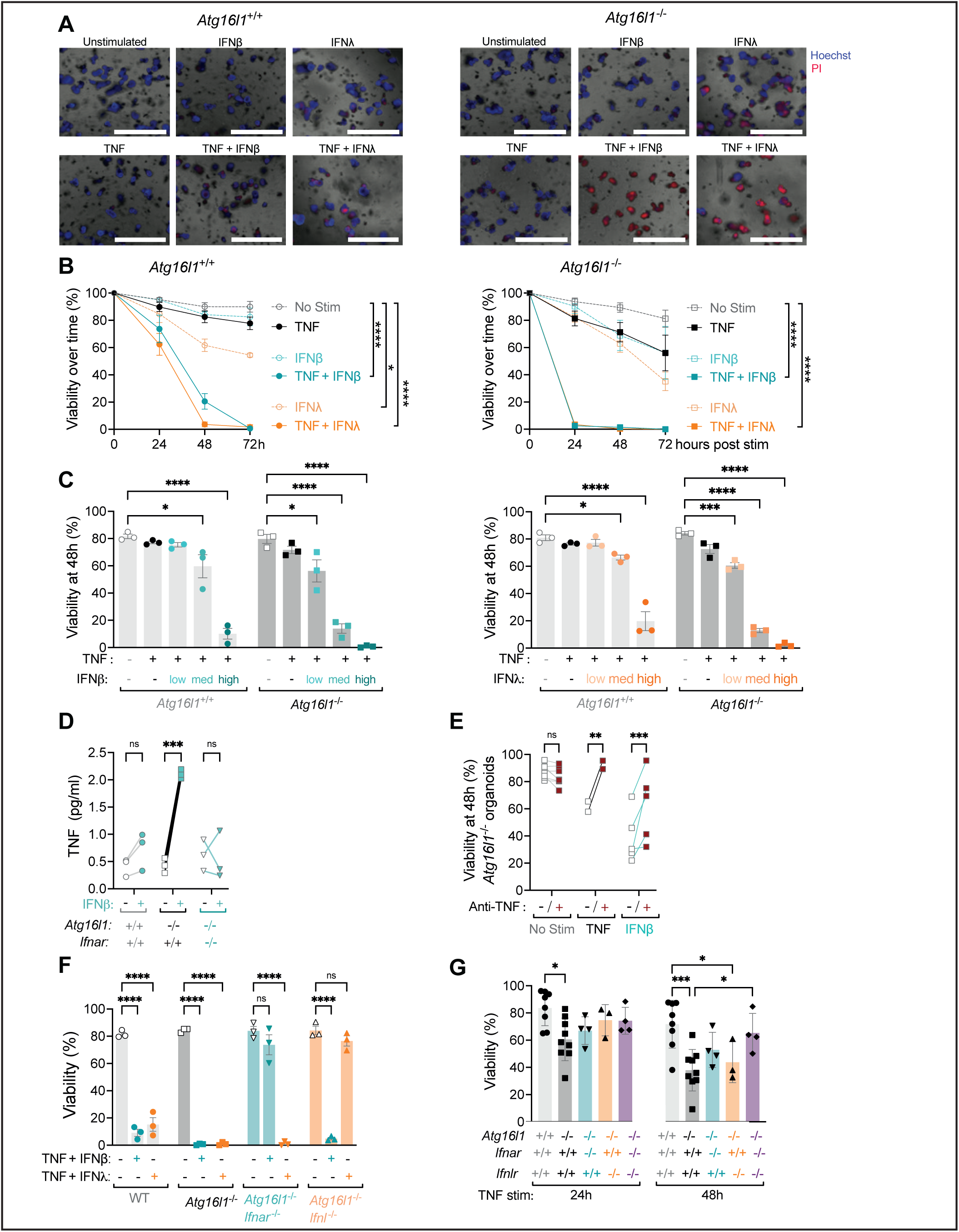
IFNβ and IFNλ2 exacerbate TNF-mediated cell death of intestinal organoids. **(A)** Representative images of small intestinal organoids derived from WT (*Atg16l1^+/+^*) and *Atg16l1^ΔIEC^* mice (*Atg16l1^−/−^*) cultivated in ENR medium for 3 days and stimulated for another 24h ± 20 ng/ml TNF, ± 100 U/ml IFNβ and ± 1ng/ml IFNλ2, then stained with Hoechst (blue) for nuclei and propidium iodide (PI) (red) for cell death. Scale bars represent 1000 μm. **(B)** Viability of *Atg16l1^+/+^* and *Atg16l1^−/−^* organoids stimulated with indicated cytokines over time as determined by PI staining and organoid morphology. **(C)** Viability of organoids 48h post stimulation with 20 ng/ml TNF and 1 (low), 10 (med), 100 (high) U/ml IFNβ (left panel) or 0.01 (low), 0.1 (med), 1 (high) ng/ml IFNλ2 (right panel). **(D)** Quantification of TNF in culture supernatants by ELISA from indicated organoids stimulated or not with 100 U/ml IFNβ for 48h. **(E)** Viability of *Atg16l1*^+/+^ (left) or *Atg16l1*^−/−^ (right) organoids treated with 20 µg/ml anti-TNFα antibody or isotype control for at least 30min before stimulation with cytokines. **(F)** Viability of *Atg16l1^+/+^*, *Atg16l1^−/−^*, *Atg16l1^−/−^Ifnar^−/−^* (derived from *Atg16l1;Ifnar*^ΔIEC^ mice), and *Atg16l1^−/−^Ifnlr^−/−^* organoids (derived from *Atg16l1;Ifnlr*^ΔIEC^ mice) stimulated on day 3 for 48h. **(G)** Viability of organoids from the indicated genotype stimulated 2h post seeding with 20 ng/ml TNF for 24 and 48h. For each condition, at least 20 organoids were seeded in 3 different wells (technical triplicate), which was repeated in at least 3 independent experiments with organoids derived from different mice each repeat. Each symbol represents the average of one technical triplicate (or the mean for each stimulation in B) with bars representing mean ± SEM. ns = not significant, *p≤0.05, **p≤0.01, ****p≤0.0001. See also Figure S2.

IFN-β can induce TNF production^13^, which may explain why IFN treatment alone can induce some degree of cytotoxicity to organoids (Fig. 2B). We observed that *Tnf* gene expression and protein secretion were increased in *Atg16l1^−/−^*organoids but not *Atg16L1^+/+^* organoids treated with IFN-β (Fig. 2D and Sup. Fig. 2A). Antibody-mediated blockade of TNF rescued viability of *Atg16l1^−/−^*organoids stimulated with TNF as expected and also greatly improved viability of *Atg16l1^−/−^* organoids treated with IFN-β (Fig. 2E). Anti-TNF improved viability of *Atg16L1^+/+^*and *Atg16l1^−/−^* organoids treated with TNF + IFN-λ2 but not IFN-λ2 alone (Sup. Fig. 2B). These results indicate that IFN-β increases the production of TNF in *Atg16l1^−/−^*organoids, which contributes to its toxicity.

We confirmed the specificity of IFN-β and IFN-λ2 for their cognate receptors. *Atg16l1^−/−^ Ifnar^−/−^* organoids were resistant to TNF + IFN-β and *Atg16l1^−/−^Ifnlr ^−/−^* organoids were resistant to TNF + IFN-λ2 (Fig. 2F). Also, IFN-β and IFN-λ2 did not induce representative ISGs in the absence of their respective receptors (Sup. Fig. 2C). However, the spontaneous ISG expression observed in untreated *Atg16l1^−/−^* organoids was absent in *Atg16l1^−/−^Ifnlr ^−/−^* but not *Atg16l1^−/−^Ifnar^−/−^*organoids (Sup. Fig. 2C), suggesting a dominant role for IFNLR, possibly due to the overexpression of *Ifnl2* in absence of IFNAR as previously described^47^. The necroptosis gene *Mlkl* is also an ISG and displayed similar IFNAR and IFNLR dependent expression patterns (Sup. Fig. 2C). Given our prior finding that ATG16L1-deficient organoids display spontaneous ISG expression^10^, we determined whether IFN receptor signaling contributed to cell death induced by TNF alone. For these experiments, we added TNF to organoids on day 0, a condition that allows us to examine toxicity of TNF treatment alone. Although the effect of deleting IFNAR or IFNLR individually (*Atg16l1^−/−^Ifnar^−/−^*and *Atg16l1^−/−^Ifnlr^−/−^*) was modest, deleting both IFN receptors improved the viability of *Atg16l1^−/−^* organoids (*Atg16l1^−/−^Ifnar^−/−^Ifnlr^−/−^*) (Fig. 2G). When these experiments using exogenous cytokines and receptor deletions are taken together, they show that IFNs synergize with TNF to mediate cell death, especially in the ATG16L1-deficient setting.

### Exacerbation of TNF-induced toxicity in organoids by IFNs is partially dependent on necroptosis signaling

TNF receptor engagement in IECs leads to activation of receptor-interacting serine/threonine protein kinase 1 (RIPK1) to mediate either apoptosis or necroptosis in IECs^48,49^. RIPK1 mediates apoptosis through caspases, and when apoptosis or caspases are inhibited, necroptosis occurs through forming a complex with RIPK3 to phosphorylate the pore-forming molecule MLKL that disrupts cell membrane integrity. We and others have shown that TNF-induced cell death in *Atg16l1* deficient intestinal organoids and macrophages is dependent on RIPK1, RIPK3, and MLKL^7, 13, 50^. To investigate how the combination of TNF and IFN reduced viability, we treated organoids with ruxolitinib as a control, a pan-caspase inhibitor (QVD), and a RIPK1-inhibitor (Nec1s). Ruxolitinib and Nec1s improved viability of both WT and *Atg16l1*-deficient organoids treated with TNF + IFN-λ2 and TNF + IFN-β (Fig. 3A, B, Sup. Fig. 3A). By contrast, QVD did not improve the viability of TNF + IFN stimulated organoids.

**Figure 3:**
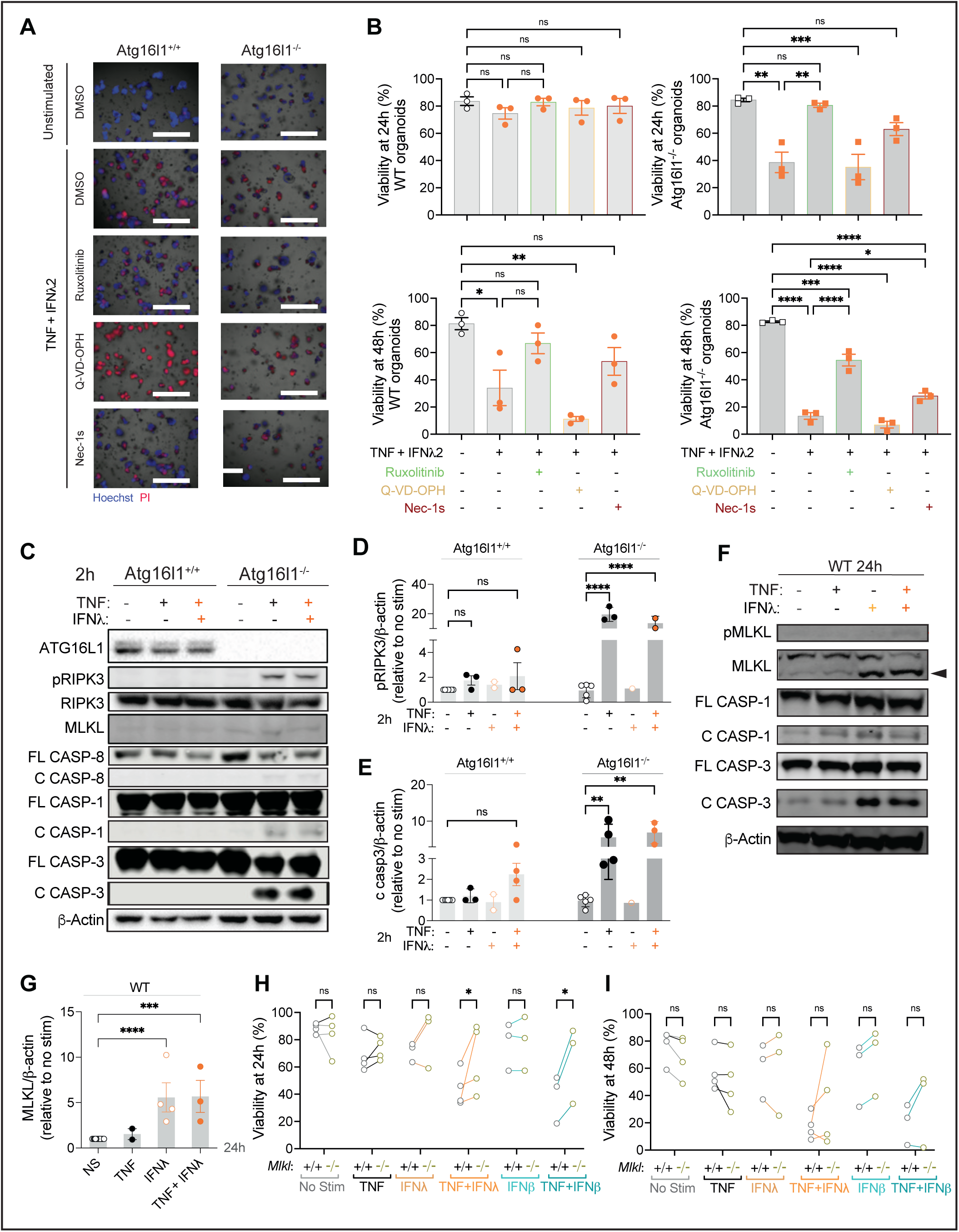
TNF and IFNλ synergize to promote RIPK1-dependent cell death in small intestinal organoids. **(A)** Representative images of Hoechst (blue) and PI (red) staining of small intestinal organoids from WT (*Atg16l1^+/+^*) and *Atg16l1^ΔIEC^* mice (*Atg16l1^−/−^*) cultivated in ENR medium for 3 days and stimulated with ± 20 ng/ml TNF, ± 1ng/ml IFNλ2 for 48h. Organoids were pre-treated with vehicle control, 100nM Ruxolitinib, 10μM Q-VD-OPh, or 10 µM Nec-1s for at least 30min before TNF+IFNλ2 stimulation. Scale bars represent 1000µm. **(B)** Quantification of organoid viability from (A) at 24 and 48h post-stimulation. For each condition, at least 20 organoids were seeded in 3 different wells (technical triplicate), which was repeated in 3 independent experiments with organoids derived from different mice). **(C-E)** Representative western blot of indicated proteins (C) and quantification of phosphorylated RIPK3 (pRIPK3) (D) and cleaved caspase-3 normalized to β-actin and unstimulated control (E) in lysates from *Atg16l1^+/+^* and *Atg16l1^−/−^*organoids stimulated for 2h with 20 ng/ml TNF alone or combined with 1ng/ml IFNλ2. C and FL designate cleaved and full length, respectively. **(F-G)** Representative western blot of indicated proteins (F) and quantification of MLKL normalized to β-actin and unstimulated control (G) in lysates from *Atg16l1^+/+^* organoids stimulated for 24h with 20 ng/ml TNF alone, 1ng/ml IFNλ2 alone or the combination of both. **(H-I)** Viability of organoids derived from WT and *Mlkl^−/−^* mice treated at day 3 with the indicated combinations of TNF, IFNλ2, and IFNβ for 24h (H) or 48h (I). Each symbol represents the average of one technical triplicate (B, I-J) or protein expression observed from a pool of different wells in an independent experiment (F-H) with bars representing mean ± SEM. ns = not significant, *p≤0.05, **p≤0.01, ****p≤0.0001. See also Figure S2 and S3.

We first examined protein levels of cell death mediators at an early time point of cytokine treatment before *Atg16l1*^−/−^ organoids displayed signs of cytotoxicity. Even at 2 h post-stimulation, phosphorylated RIPK3 (pRIPK3) and cleaved caspases, especially caspase-3, were visible in TNF or TNF + IFN stimulated *Atg16l1*^−/−^ organoids and not WT organoids (Fig. 3C-F, Sup. Fig. 3B). This result is consistent with the delayed onset of cell death in WT organoids. MLKL and cleaved caspase-3 were detected in WT organoids treated with IFN-λ2 for 24 h, with or without TNF; pMLKL was faintly visible when both cytokines were present (Fig. 3F, G, Sup. Fig. 3B). Although our earlier observation implicated necroptosis because RIPK1 inhibition and not caspase inhibition protected organoids, we noted the prominent cleavage of caspase-3. Therefore, we tested whether cell death induced by TNF and IFN treatment were dependent on MLKL. *Mlkl^−/−^* organoids displayed increased viability when treated with TNF + IFN-λ2 or TNF + IFN-β at 24 h, but this protection was less consistent at 48 h (Fig. 3I, H). These results indicate that necroptosis contributes to the loss of viability induced by TNF + IFN treatment of organoids.

### Virally-triggered intestinal disease in Atg16l1 mutant mice is RIPK3-dependent

Paneth cell abnormalities and mortality are ameliorated by RIPK1 inhibition in MNV + DSS-treated *Atg16l1^ΔIEC^* mice^7^. As a kinase, RIPK3 may also be a targetable molecule. We found that *Atg16l1^ΔIEC^;Ripk3^−/−^* mice showed substantially reduced lethality and weight loss compared to *Atg16l1^ΔIEC^*mice following MNV + DSS treatment similar to WT and *Ripk3^−/−^*control mice (Fig. 4A). Although not a complete reversal, disease score and colon length of *Atg16l1^ΔIEC^;Ripk3^−/−^*mice were not significantly different from WT and *Ripk3^−/−^* mice (Fig. 4A, B), indicating a general improvement in disease outcomes. Additionally, Paneth cell numbers and lysozyme staining were restored in *Atg16l1^ΔIEC^;Ripk3^−/−^* mice (Fig. 4C-E). These results show that RIPK3 is necessary for susceptibility of *Atg16l1^ΔIEC^*mice to MNV.

**Figure 4:**
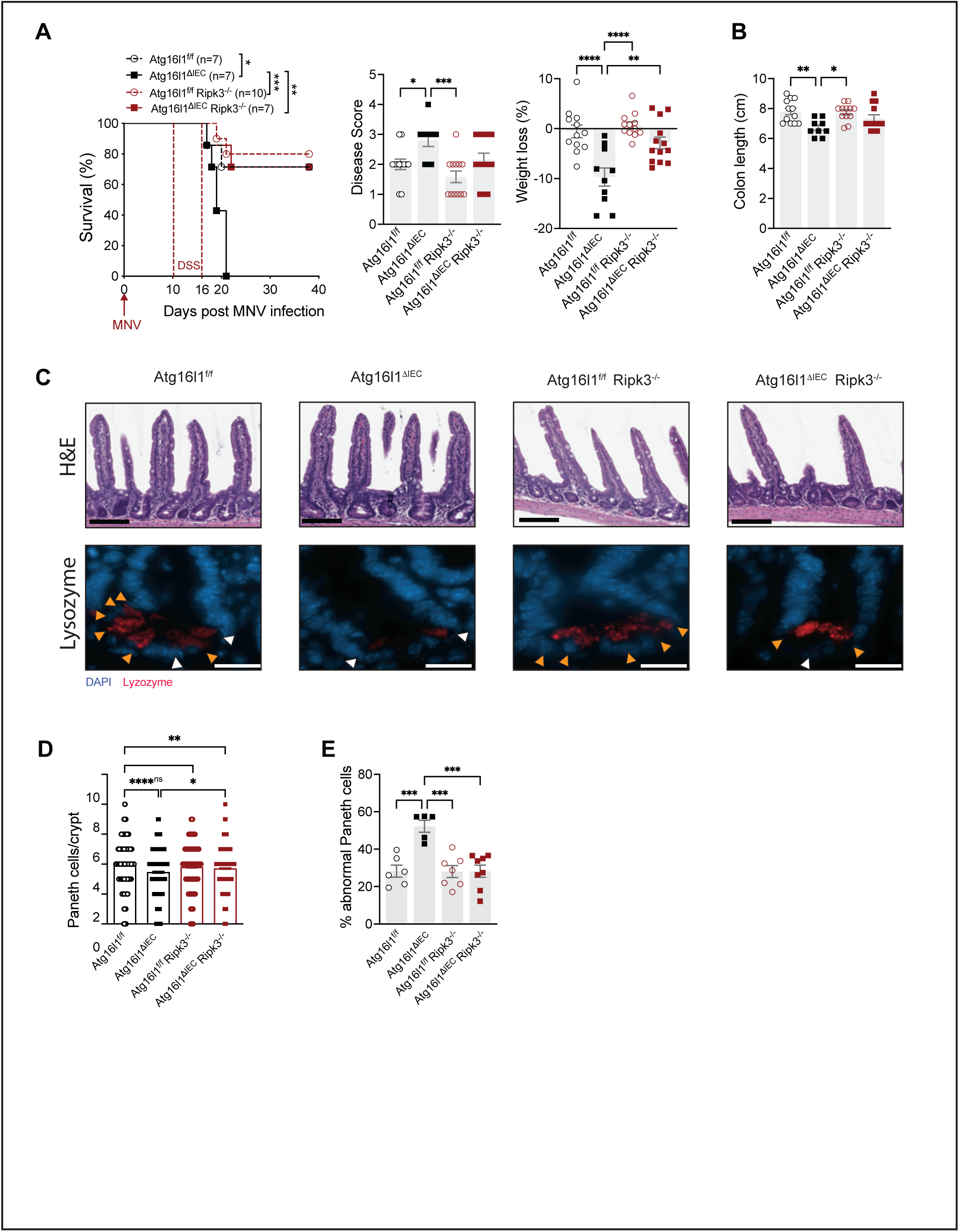
Virally-triggered intestinal disease in Atg16l1 mutant mice is RIPK3-dependent. **(A)** Survival over time, cumulative disease score, and weight loss in *Atg16L1*^f/f^, *Atg16L1*^ΔIEC^, *Atg16L1*^f/f^;*Ripk*3^−/−^, and *Atg16L1*^ΔIEC^;*Ripk3*^−/−^ mice infected for 10 days with MNV before receiving 3% DSS in their drinking water for 6 days. **(B)** Colon length on day 5 post DSS treatment of indicated mice infected with MNV as in (A). **(C)** Representative images of H&E staining (top, scale bar 100μm) and immunofluorescence of ileal crypts from mice in (B) stained for lysozyme (red) and DAPI (blue) (bottom, scale bar 20μm). **(D)** Number of Paneth cells observed in each crypt in mice from each genotype. **(E)** Average proportion of abnormal Paneth cells per crypt observed in each mouse as determined on the basis of whether each Paneth cell displayed a typical staining pattern with distinguishable granules (normal, orange arrow) or depleted and/or diffuse staining (abnormal, white arrow). At least 50 crypts were quantified per mouse. Each symbol represents individual mice with bars representing mean ± SEM from at least two independent experiments (except for Paneth cell abnormality, n=1). ns = not significant, *p≤0.05, **p≤0.01, ***p≤0.001, ****p≤0.0001.

### Human organoids are susceptible to cell death induced by antiviral factors

The combination of TNF and IFNs is toxic to colonic organoids derived from humans, although the concentration of cytokines necessary for cell death is high^51^. Based on our data with mouse organoids, we hypothesized that human small intestinal organoids will be sensitive to TNF + IFN treatment, especially in the presence of the *ATG16L1* risk allele. To test this hypothesis, we utilized organoids that we previously generated from endoscopic biopsy specimens that were genotyped for the *ATG16L1^T300A^* variant (Sup Table 1)^10^. As in the mouse model, the combination of TNF + IFN-λ2 induced cell death in human organoids, which was exacerbated in those derived from individuals homozygous for *ATG16L1^T300A^* (Fig. 5A-C).

**Figure 5:**
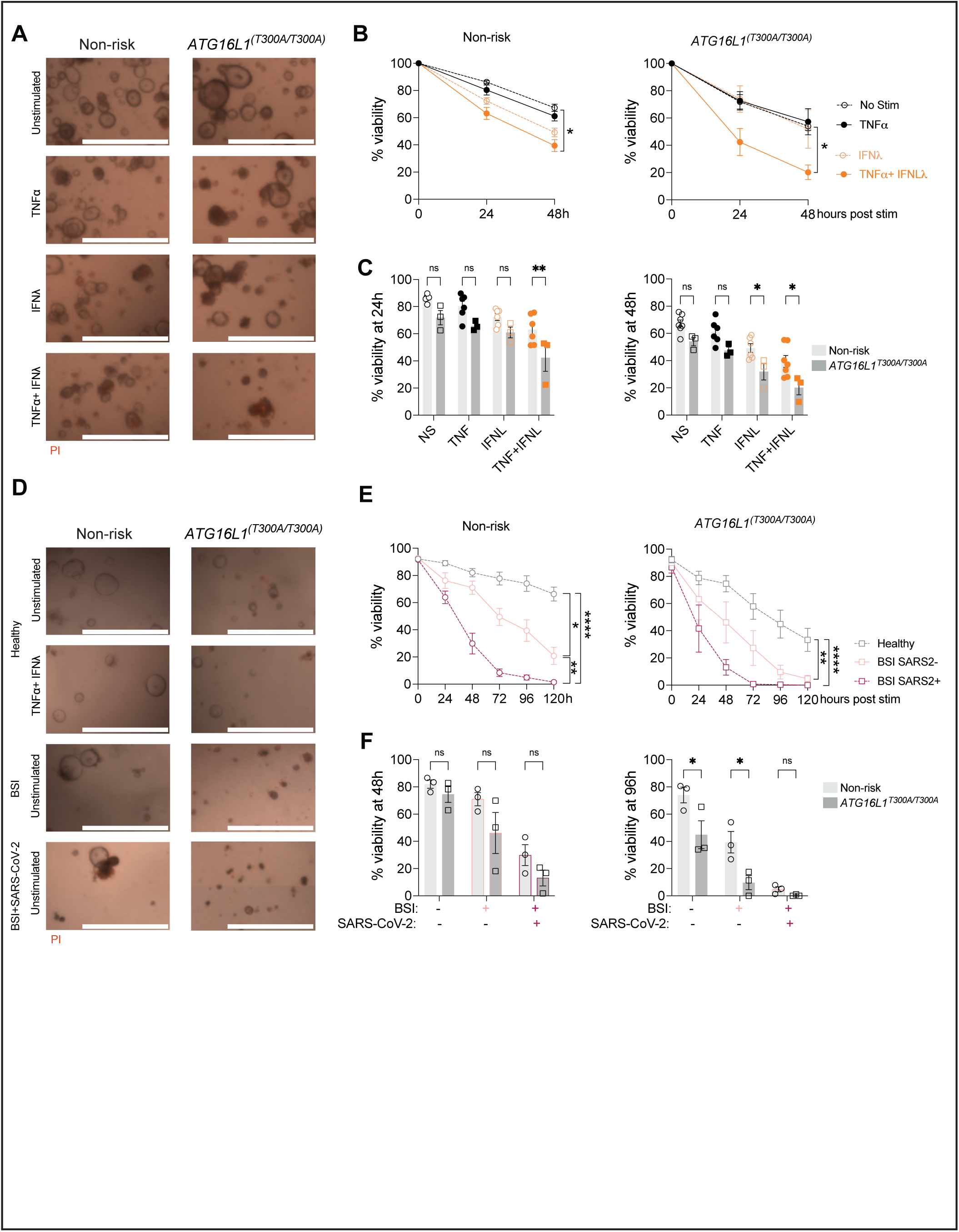
Intestinal organoids derived from *ATG16L1T300A* homozygous individuals exhibit heightened susceptibility to inflammatory factors. **(A-B)** Representative images at 24h post-stimulation with ± 50 ng/mL hTNFα, ± 100 ng/mL hIFNλ2 (A) and viability over time (B) of human organoids from controls (n = 6 donors homozygous or heterozygous for the non-risk allele) and *ATG16L1^T300A^*homozygous individuals (n=3 donors). **(C)** Bar graph representation of data from (B) comparing non-risk control and *ATG16L1^T300A^* homozygous organoids. **(D-E)** Representative pictures at 48h (D) and viability over time (E) of non-risk (n=3) and *ATG16L1T300A* homozygous (n=3) organoids post incubation with pooled serum from healthy donors, patients with a blood stream infection (BSI+ SARS-CoV-2-) or patients with a BSI and infected with SARS-CoV-2 (BSI+ SARS-CoV-2+). **(F)** Bar graph representation of data from (E) comparing non-risk control and *ATG16L1^T300A^* homozygous organoids. For each condition, at least 20 organoids were seeded in 3 different wells (technical triplicate). Each symbol represents the average viability per donor (calculated from technical triplicates) with bars representing mean ± SEM for each group. For (A-C) 2-5 independent experiments were performed for each organoid donor. For (D-E), one experiment was performed for each organoid donor due to the limited quantity of serum available. ns = not significant, *p≤0.05, **p≤0.01, ****p≤0.0001.

TNF and IFNs are produced in response to most viral infections. We previously showed that SARS-CoV-2 infected mice display a reduction in Paneth cell numbers and abnormal lysozyme staining, reminiscent of *Atg16l1* mutant mice infected with MNV^52^. We did not detect replication competent SARS-CoV-2 particles in the gut of these mice, raising the possibility that circulating antiviral cytokines can mediate long-range cytotoxic effects on IECs in situations in which they reach high levels, such as observed in severe COVID^53, 54^. Indeed, gastrointestinal symptoms are common in patients with COVID-19 and many hospitalized patients display blood stream infections (BSI) caused by microbes originating from the gut, suggesting intestinal barrier defects^52^. To test whether soluble immune mediators associated with severe COVID-19 are cytotoxic to human IECs, we incubated the above organoids with serum samples pooled from hospitalized patients during the height of the pandemic when individuals were screened for SARS-CoV-2 infection. These samples were compared with serum from healthy volunteers and hospitalized individuals who were COVID-negative but BSI-positive. Compared with serum from healthy volunteers, addition of serum from patients with severe COVID led to increased loss of viability, which was worse in organoids homozygous for *ATG16L1^T300A^* (Fig. 5D-F). Serum from BSI-positive control also worsened organoid death but not as severely as the COVID-positive samples (Fig. 5 D-F). Therefore, circulating factors associated with severe COVID-19 display cytotoxicity to IECs, and the *ATG16L1* risk variant further sensitizes cells to this activity.

## Discussion

Our study demonstrates that type I and III IFNs are central mediators of intestinal epithelial damage following viral infection in the context of ATG16L1 deficiency. Antiviral IFNs and ISG expression have been observed in IBD patients but their contribution to the pathology is incompletely understood^34, 55–57^. We show that IFN signaling in IECs drives epithelial cell death and disease exacerbation in MNV-infected *Atg16l1^ΔIEC^* mice, while organoid models reveal a synergistic cytotoxicity between IFNs and TNF associated with necroptosis signaling. Notably, deletion of IFN receptors rescued overall survival but had partial effects on Paneth cells. This may reflect a confounding role for IFNs in controlling the virus, or alternatively, redundancy with other cytokines such as IFN-γ, which is known to be upregulated during MNV infection and can also trigger Paneth cell dysfunction^8, 58, 59^. IFN-γ and type I/III IFNs display redundancy in a model of chronic murine astrovirus infection^60^. Also, the role of IFNAR was unexpected given the dominance of IFNLR signaling in IECs and may reflect selective functions in a subset of IECs or a role specific to the *Atg16l1* mutant setting. Future studies should explore whether each IFN has a distinct IEC target and timing of activity that controls their cytotoxic effects.

These findings strengthen the link between infectious agents such as noroviruses and CD^4, 61, 62^. Although noroviruses are associated with CD in some patients, they most often cause self-resolving gastroenteritis and are detected in many asymptomatic individuals, suggesting that as in our mouse model, the consequences of their presence depend on host factors and may be immune mediated^63, 64^. A recent study showed that Epstein-Bar virus (EBV) infection precedes CD^65^. Thus, triggers of intestinal inflammation are unlikely to be exclusive to noroviruses but there may be some specificity. Although many types of gut microbes including commensal bacteria evoke TNF and IFNs, our finding that serum from patients with severe COVID-19 displays higher toxicity to organoids compared with serum from patients with non-viral BSIs raises the possibility that certain types of infections are more likely to trigger responses that cause IEC death. These observations also raise the intriguing possibility that viral sequelae and chronic diseases like IBD share underlying mechanisms that include IEC death. However, we acknowledge that our experimental system does not allow for such comparisons, and that IBD is much more complex with a prolonged disease development and cycles of healing and recurrence.

Therapeutically targeting signaling molecules upstream of cell death may offer a strategy to protect the epithelium in at-risk individuals. Although RIPK3 deletion ameliorated disease in *Atg16l1^ΔIEC^*mice *in vivo*, RIPK1 inhibition may provide broader protection by blocking multiple death pathways. Crosstalk between necroptosis, apoptosis, and pyroptosis and their co-occurrence^66–68^ may be especially relevant to organoids that harbor multiple IEC types. Our time course experiments show that necroptosis inhibition delays but does not completely prevent loss of viability in organoids. The partial rescue of viability is reminiscent of findings showing that deleting *Tnfaip3* and *Tnip1* (genes that are also linked to IBD and autoimmunity) sensitizes to TNF toxicity in the presence of microbial and viral ligands^69, 70^. Interventions that can be personalized to restore function of a cell type that is specifically affected in a given susceptible host, such as Paneth cells in the ATG16L1-deficient background, may be ideal. In this sense, our observation that RIPK3 deletion better restores Paneth cells compared with IFN receptor deletion supports targeting nodes at the intersection of multiple cytokine signaling pathways. Indeed, clinically used JAK inhibitors effectively protect patient-derived colonic organoids from cytokine cocktails^51^.

In summary, this work positions IFNs as key players in virally triggered intestinal pathology associated with TNF toxicity, particularly in the setting of ATG16L1 deficiency. It underscores the need to consider synergistic cytokine responses in the management of CD and other barrier-associated disorders.

## Methods

### Mice

Age- and gender-matched 8–12-week-old mice on the C57BL/6J (B6) background were used. *Atg16l1^f/f^*;VillinCre (*Atg16l1^ΔIEC^*) mice were provided by S. Virgin (Washington University School of Medicine, St. Louis, MO) and crossed to *Ifnar1^f/f^* (Jax, 028256) and *Ifnlr1^f/f^* mice (provided by S. Virgin) to obtain *Atg16l1;Ifnar^ΔIEC^* and *Atg16l1;Ifnlr^ΔIEC^*mice, respectively. To generate littermate VillinCre-negative and *Atg16l1^ΔIEC^*controls, mice were bred heterozygous for either the *Ifnar^f^* or *Ifnlr^f^* genes and crossed VillinCre-positive with VillinCre-negative. Ifnar*^ΔIEC^*and *Ifnlr^ΔIEC^* mice were bred separately by crossing VillinCre-positive with VillinCre-negative mice. *Atg16l1;Ifnlr;Ifnar^ΔIEC^*mice were obtained by crossing *Atg16l1;Ifnar1^ΔIEC^* and *Atg16l1;Ifnlr^ΔIEC^*mice. WT mice are a pool of VillinCre-negative mice from different crosses. *Rip3*^−/−^ mice were provided by G. Miller (NYU School of Medicine, New York, NY). *Mlkl^−/−^* mice were provided by J. Murphy (WEHI, Parkville, Australia). All animal studies were performed according to approved protocols by the New York University School of Medicine Institutional Animal Care and Use Committees.

### MNV-triggered disease model

MNV.CR6 stock was prepared as previously described and as detailed in the Supplementary method section^71^. Mice were infected orally with 3 × 10^6^ PFUs resuspended in PBS. Viral burden in stool at day 10 post infection was determined by plaque assay. On day 10 post-infection, mice were administered 5% Dextran Sulfate Sodium (DSS) (TdB Consultancy) in their drinking water for 6 d after which it was replaced by regular drinking water. *Ripk3*^−/−^ mice showed increased lethality when given 5% DSS. When these mice were used, 3% DSS was used instead of 5% as indicated in the figure legend. Clinical disease score and mouse weight were assessed daily. Disease score was quantified on the basis of five parameters as described previously, including diarrhea (0–2), hunched posture (0–2), fur ruffling (0 or 1), mobility (0–2), and blood stool (0 or 1), in which 8 was the maximum score for the pathology^7^. For survival experiments, mice were euthanized if they reached the humane end point. In other experiments, mice were sacrificed before receiving DSS or after receiving 5 days of DSS (which occurred before clinical disease signs).

### Intestinal organoid culture

Murine and human organoids (generated during previous studies^10, 72^) were cultured as described in the Supplementary methods. Murine organoids were treated with 20 ng/ml mTNFα (Peprotech) and indicated concentrations of mIFNβ (PBL assay science) and mIFNλ2 (Peprotech). In some experiments, organoids were pre-treated with 100nM Ruxolitinib, 10μM Q-VD-OPh (Millipore Sigma), 10 µM Necrostatin-1s (Nec-1s) (Biovision), 20 µg/ml anti-TNFα antibody (XT3.11, BioXcell) or 20 µg/ml Rat IgG1 isotype control (HRPN, BioXcell). Human organoids were treated with 50 ng/mL hTNFα (Peprotech), 100 ng/mL hIFNλ2 (R&D systems). In another experiment, half of the DMEM/F-12 in the differentiation medium was replaced by human serum (growth factors were maintained at same concentrations). Serum from healthy donors from the United States was purchased from SeraCare. The serum from patients with BSI and SARS-CoV-2 infection was collected at the NYU Langone hospital in 2020 during a previous study (Table S3)^73^. Each organoid donor was stimulated with the same pool of serum obtained from healthy donors, patients with BSI and negative for SARS-CoV-2 by qPCR or with BSI and positive for SARS-CoV-2. Percentages of viable organoids were determined based on morphology by daily quantification as before^7^. Dead organoids were also marked by staining with 100 µg/ml PI (Sigma-Aldrich) and 100 µg/ml Hoechst 33342 (Invitrogen) to confirm phenotypes.

### Statistical analysis

GraphPad Prism version 10 was used for statistical analysis unless otherwise specified. Survival data were analyzed by Kaplan-Meier with Log-rank test followed by Holm Sidak’s multiple comparison test (Figure 1A, 4A). For other parameters, ordinary or lognormal ordinary (PFU, Figure 1D) one-way ANOVA and Tukey’s multiple comparison were used when data followed a normal distribution or a Kruskal-Wallis test followed by Dunn’s multiple comparison test when they did not (disease score, Figure 1B). For histological measurements, the effect of mouse genotype was determined on R using individual measurements (e.g. number of Paneth cells in 50 crypts per mouse) while adjusting for batch effect caused by individual mouse and experiments. ANOVA type III followed by Tukey’s multiple comparison Generalized Linear Mixed Model (GLMM) was used for Paneth cell counts. A beta-binomial analysis followed by Tukey’s multiple comparison was used for Paneth cell abnormality. Negative Binomial GLMM was used to compare the number of TUNEL+ cells per crypt. In Figure 2 and 3, in each experiment, organoids derived from a single mouse per genotype were stimulated with different cytokines and inhibitors (experiments were then repeated in independent experiments using organoids derived from different mice). Ordinary one-way ANOVA were used to compare the effect of the treatment within a genotype, two-way ANOVA were used when comparing the effect of the treatment and of the genotypes, followed by Sidak’s multiple comparison test. Comparison of area under the curve was performed to compare time-course curves. For human organoids, an ANOVA followed by Sidak’s multiple comparison test was used to compare the effect of the stimulation on organoid viability depending on the genotype.

## Data availability

All data generated or analysed during this study are included in this published article and its supplementary information files.

## Acknowledgements

We thank the NYU Langone’s Experimental Pathology Research Laboratory (RRID:SCR_017928) and the NYU Langone’s Microscopy Laboratory (RRID: SCR_017934), supported in part by NYU Langone Health’s Laura and Isaac Perlmutter Cancer Center Support (grant P30CA016087), for the use of their instruments and technical assistance. We thank the following University of Pennsylvania core facilities for their expert technical support: the Cell and Developmental Biology (CDB) Microscopy Core (RRID SCR_022373) as well as the Molecular Pathology & Imaging Core (MPIC) (RRID: SCR_022420) and Genetically Engineered Mouse Core supported by the UPENN Digestive and Liver Center (P30DK050306). We also thank M. B. da Silva, Y. Shono and M. R. M. van den Brink for their support with TUNEL staining and N Desvignes and A Lambert for their help with lysozyme staining. We also thank Filipe De Vadder for his kind help with statistical analyses. This research was funded in part by NIH grants DK093668 (K.C.), AI121244 (K.C.), AI140754 (K.C.), AI179896 (K.C.), and DK050306 (K.C.); Shanahan Family Fund for Inflammatory Bowel Disease Research (K.C.); pilot grant from King Center for Lynch Syndrome (K.C.); pilot grant from Abramson Cancer Center (K.C.); and an industry partnership with AbbVie Inc. (K.C.).

## Author contributions

LBR, JAN, YMI and KC designed the study. LBR, JAN and YMI performed most of the experiments with assistance from KK, TH, and DM and BMM. YMI and JA provided the human organoids lines. AL, AD and VJT provided the human serum samples. LBR, JAN and KC drafted the manuscript. All authors contributed to refinement of the study protocol and approved the final manuscript.

