## Supplementary material for "Type I and III interferons synergize with TNF to promote virally-triggered damage to the intestinal epithelium"

**Supplementary methods**

*MNV stock generation*

MNV.CR6 stock was prepared as previously described[1]. In brief, supernatant from 293T cells transfected with a plasmid containing the MNV.CR6 genome was used to inoculate to RAW264.7 cells. Infectious virus in the supernatant was concentrated by ultracentrifugation and resuspended in endotoxin-free PBS. Virus titer was determined by plaque assay as previously described with few modifications [2]. Diluted virus was incubated on confluent RAW264.7 cells for 1h followed by overlay with 10g/L methylcellulose media (Sigma-Aldrich). Cells were incubated at 37°C for 72 h and stained with 1g/L crystal violet solution.

*Intestinal organoid culture*

Murine small intestinal crypts were isolated and cultured as described previously using ENR media composed of DMEM/F-12 (ThermoFisher) in the presence of 100IU Penicillin and 100 μg/ml Streptomycin (Corning), 125 μg/ml Gentamicin (ThermoFisher), 2 mM-Glutamine (Corning), 20 ng/ml mEGF (PeproTech), 100 ng/ml mNoggin (R&D systems), and 500 ng/ml mR-Spondin 1 (R&D systems)[3]. Human organoids were generated during previous studies and frozen[4, 5]. For this study, they were thawed and cultured in Human IntestiCult Organoid Growth Medium (Stemcell Technologies). For viability assays, crypts were embedded at 50 crypts /10 μl Matrigel (Stemcell Technologies) and cultured in triplicate for 3 days in 96-well culture plates. Assays were performed in ENR medium for mouse organoids or in differentiation medium for human organoids. Differentiation medium was composed of DMEM/F-12 containing 100ng/mL rhIGF-1 (Biolegend), 50ng/mL hFGF-basic (PeproTech), 1μg/mL rhR-Spondin 1 (R&D systems), 100ng/mL rmWnt-3A (R&D systems), 100ng/mL rmNoggin (R&D systems), 10mM Nicotinamide (Millipore), 10 nM

[Leu15]- Gastrin 1 human (Sigma-Aldrich), 500nM A 83-01 (TOCRIS), 1X B-27 supplement (ThermoFisher) as in[6, 7].

*Quantitative qPCR*

RNA was extracted from organoids and intestinal tissue using the RNeasy mini kit (Qiagen). DNA was removed using the RNA-free DNase set (Qiagen). cDNA was synthesized using the High-Capacity cDNA reverse transcription kit (Thermo Fisher) according to the manufacturer's protocol. Quantitative PCR was performed on a Roche 480 II LightCycler using SYBR Green I Master mix (Roche). Gene expression was normalized to either *Gapdh* (organoids) or *18S* (tissue). All primers are listed in Table S2.

*Immunoblotting*

600 small intestinal organoids were seeded in 24 well plates and cultured for 3 days in ENR before cytokine stimulation with 20 ng/ml mTNFα in the presence of either 100 U/ml IFNβ or 1 ng/ml IFNλ2 for 2-24h. Organoids were incubated in lysis buffer (20 mM Tris-HCl, pH 7.4, 150 mM NaCl, 1% Triton X-100, 10% glycerol) containing Halt protease and phosphatase inhibitor cocktail (Thermo Fisher Scientific) and run on a 4%–12% Bis-Tris Plus Gel (Invitrogen) as before[7]. The following antibodies were used for immunoblotting studies: anti–β-actin (AC-15; Sigma-Aldrich), anti–Phospho-MLKL (Ser345), anti-MLKL (Cell Signaling), anti-Phospho-RIP3 (phospho S232; Abcam), anti-RIP3 (AHP1797; AbD Serotec), anti-caspase-1 (AG-20B-0042-C100, AdipoGen) anti-caspase-3 (9662, Cell Signaling), anti-caspase-8 (ALX-804-447, Enzo) and anti-Atg16L (Sigma-Aldrich). Secondary antibodies IRDye 680RD goat anti–rabbit IgG (925-68071) and IRDye 800CW goat anti–mouse IgG (925-32210) were purchased from LI-COR.

*Histology and Immunohistochemistry*

Quantification of all microscopy data were performed blind. Intestinal sections were prepared and stained as previously described[7]. Terminal deoxynucleotidyl transferase dUTP nick end labeling (TUNEL) staining was performed using the In situ Cell Death Detection POD kit (Roche Diagnostics) on the Discovery XT according to the manufacturer's protocol. Lysozyme staining was performed using anti-lysozyme (ab108508, Abcam) and DAPI immunostaining. All slides were analyzed using a Zeiss AxioObserver.Z1 with Axiocam 503 Mono operated with Zen Blue software. Images were processed and quantified using QuPath. At least 50 villi or crypts per mouse were quantified. Abnormal Paneth cells were defined as Paneth cells showing disordered, depleted and/or diffuse lysozyme staining as in[7, 8] and adapted from[9].

### Supplementary figures

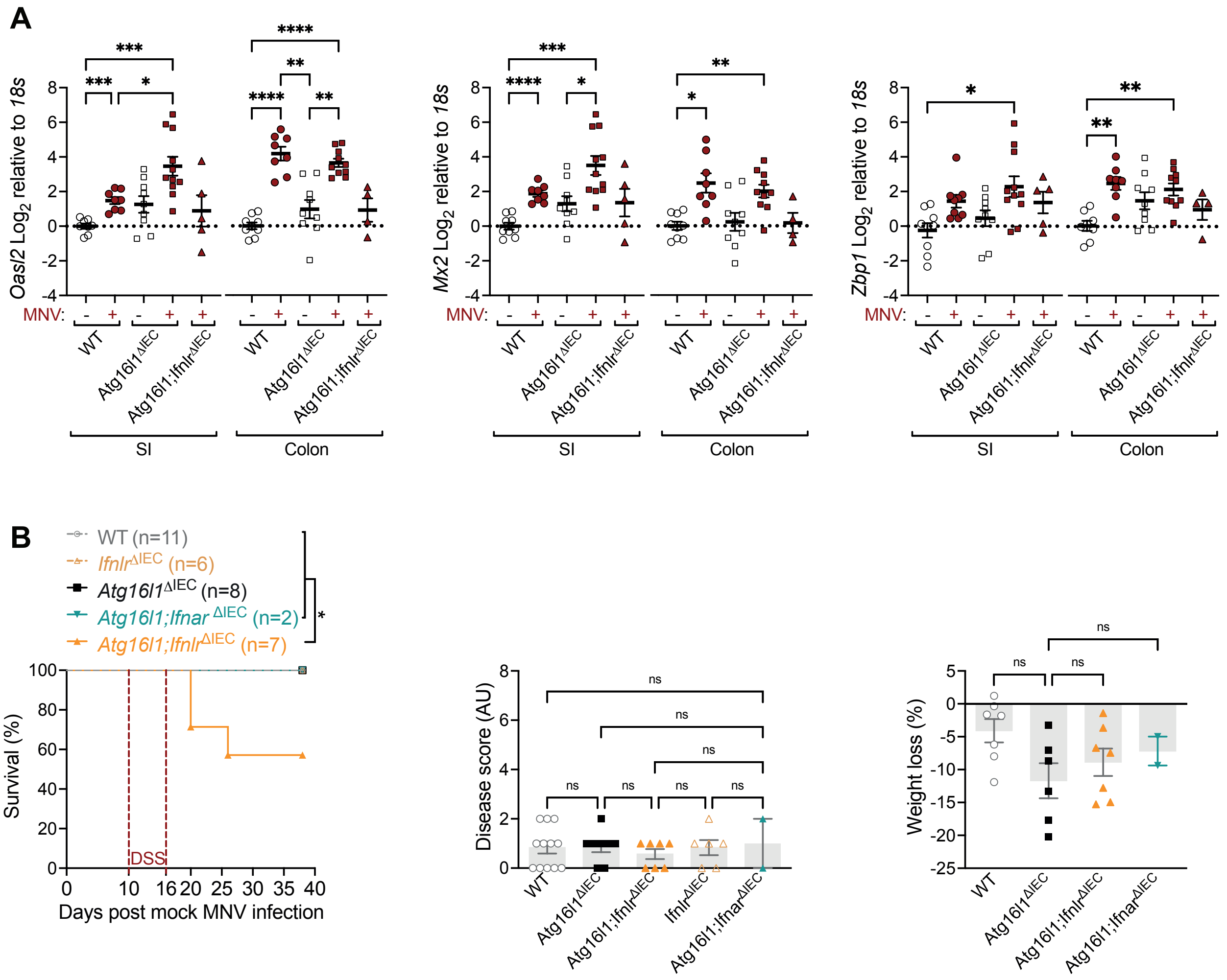


**Supplementary figure 1: Characterization of MNV infected and uninfected *Atg16L1*$^{\Delta IEC}$ mice. (A)** Normalized RNA levels of *Oasl2*, *Mx2* and *Zbp1* in the ileum (SI) or colon of WT, *Atg16l1*$^{\Delta IEC}$, and *Atg16l1;Ifnlr*$^{\Delta IEC}$ mice infected or not with MNV CR6 for 10 days. **(B)** Survival, clinical score, and weight loss of mock infected WT, *Atg16l1*$^{\Delta IEC}$, *Atg16l1;Ifnlr*$^{\Delta IEC}$ and *Atg16l1;Ifnar*$^{\Delta IEC}$ mice receiving 5% DSS in their drinking water for 6 days. Dots represents individual mice and line (A) or bars (B) represent mean ± SEM from at least two independent experiments analyzed by ordinary one-way ANOVA and Tukey's multiple comparison (area under the curves were compared for B). Related to Fig. 1.

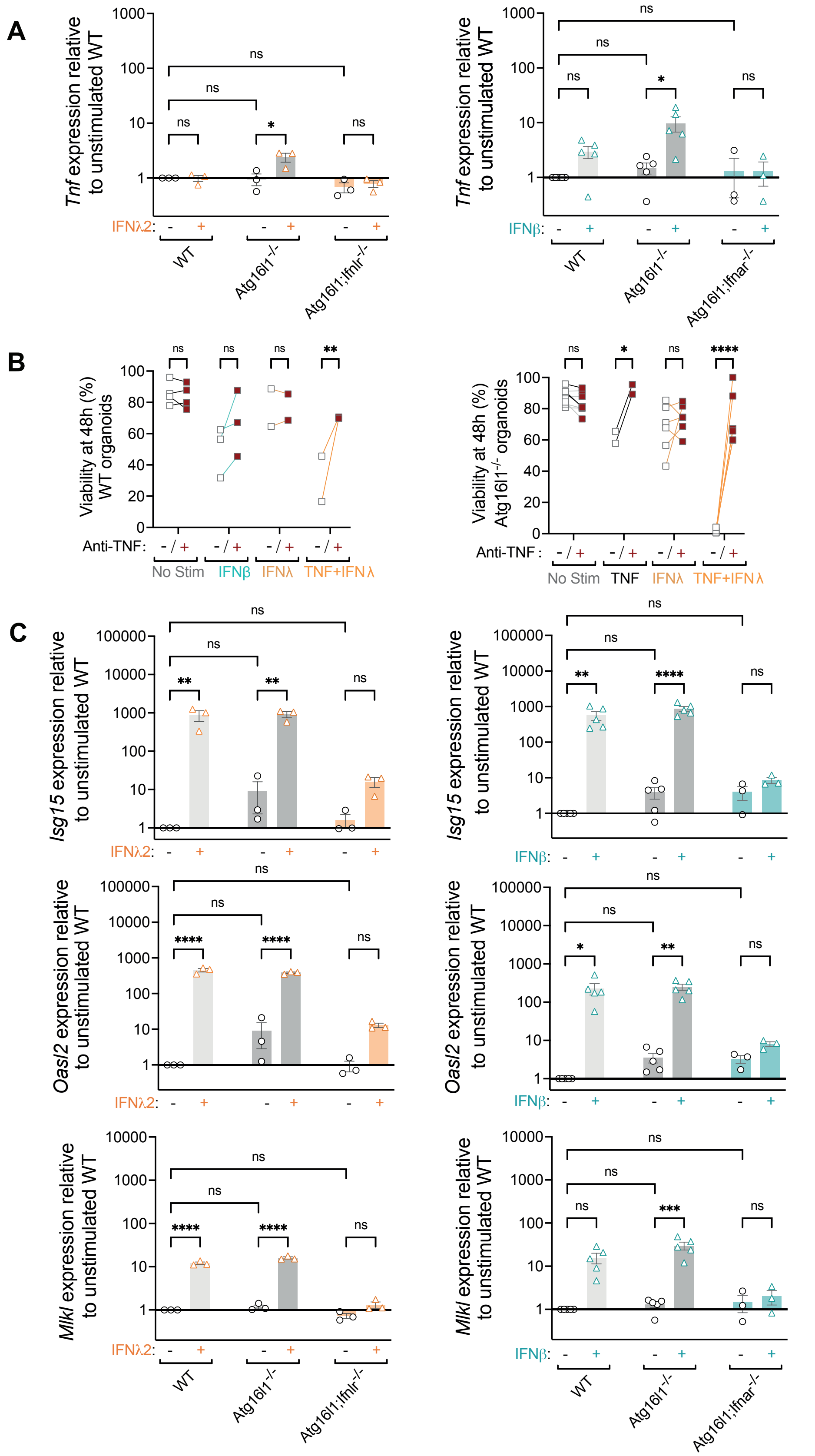
A
Tnf expression relative to unstimulated WT
IFNλ2:
IFNβ:
WT
Atg16l1-/-
Atg16l1;Ifnlr-/-
Atg16l1;Ifnar-/-
ns
*
B
Viability at 48h (%) WT organoids
Viability at 48h (%) Atg16l1-/- organoids
Anti-TNF:
- / +
No Stim
IFNβ
IFNλ
TNF+IFNλ
TNF
**
****
C
Isg15 expression relative to unstimulated WT
Oasl2 expression relative to unstimulated WT
Mlkl expression relative to unstimulated WT
***

**Supplementary figure 2: Analysis of Tnf, ISG and cell death related transcripts in murine organoids. (A)** Small intestinal organoids derived from WT, *Atg16l1*$^{\Delta IEC}$, *Atg16l1;Ifnar*$^{\Delta IEC}$, or Atg16l1;Ifnlr$^{\Delta IEC}$ mice were cultivated in ENR medium for 3 days before stimulation with ± 100 U/ml IFNβ or ± 1ng/ml IFNλ2 for 24h. Normalized RNA levels to GAPDH and WT unstimulated organoids are depicted. RNA levels were measured on an average of 600 organoids derived from at least 2 different mice in independent experiments for each condition (as depicted by each symbol). **(B)** Viability of *Atg16l1*$^{+/+}$ (left) or *Atg16l1*$^{-/-}$ (right) organoids treated with 20 µg/ml anti-TNF antibody or isotype control for at least 30min before stimulation with cytokines. **(C)** Normalized RNA levels of ISGs as measured in (A). Bars represent mean ± SEM from at least two independent experiments analyzed by ordinary one-way ANOVA and Tukey's multiple comparison. Relative to Fig. 2 and 3.

A

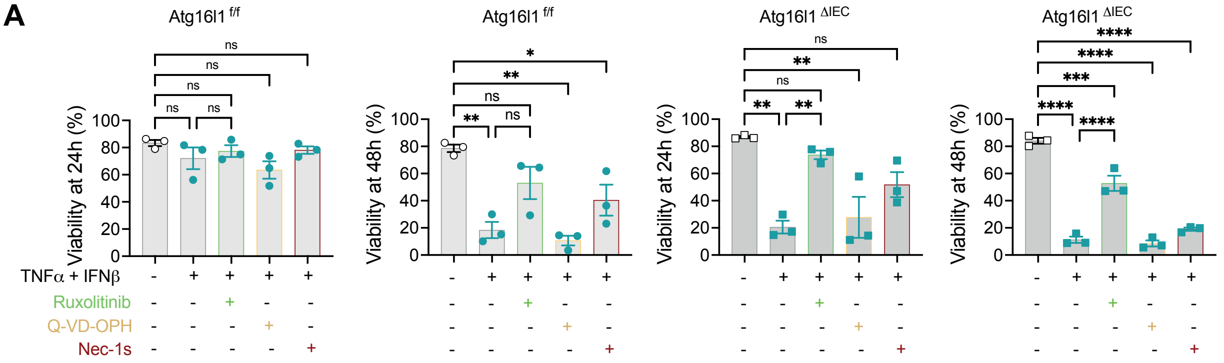


B

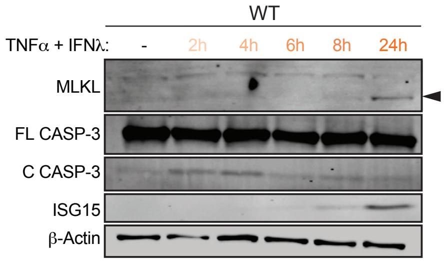


**Supplementary figure 3: IFNβ and IFNλ2 contribute to necroptosis and other cell death in small intestinal organoids.** Small intestinal organoids were cultivated in ENR medium for 3 days before stimulation with ± 20 ng/ml TNFα ±100U/ml IFNβ,. Viability is determined on the basis of PI staining and morphology before stimulation and every 24h. **(A)** Viability of WT or *Atg16l1*$^{\Delta IEC}$ organoids pre-treated with 100nM Ruxolitinib, 10μM Q-VD-OPh, or 10 μM Nec-1s for at least 30min before TNFα+IFNβ stimulation shown after 24h and 48h. Each symbol represents the average viability observed for each condition on a technical triplicate done in an independent experiment with bars representing mean ± SEM. **(B)** Representative Western blot for analysis of cell death–related proteins. Protein lysate from approximately 600 organoids stimulated for 2-24h with the indicated cytokines was used for each condition. ns = not significant by ordinary one-way ANOVA and Tukey's multiple comparison. Relative to Fig. 3 and 4.

**Table S1: Patient information for human specimens used to derive organoids**

| N° | ATG16L1$^{T300A}$ | Inflammation | Tissue | Sex | Age | Figures | Original study |
|---|---|---|---|---|---|---|---|
| **1** | (T;T) | non-inflamed | duodenum | F | 34 | 5A-C, E, F | |
| **2** | (T;A) | non-inflamed | duodenum | M | 23 | 5B, C, E, F | |
| **3** | (T;A) | non-inflamed | duodenum | M | 47 | 5B, C | |
| **4** | (T;A) | inflammed | ileum | F | 22 | 5B, C | [4] |
| **5** | (T;A) | non-inflamed | duodenum | M | 37 | 5B, C | |
| **6** | (T;A) | non-inflamed | duodenum | F | 32 | 5B, C | |
| **7** | (T;A) | non-inflamed | duodenum | F | 43 | 5B-E | |
| **8** | (T;A) | non-inflamed | ascending colon | M | 22 | 5B, C | |
| **9** | (A;A) | non-inflamed | duodenum | M | 21 | 5A-E | [4] |
| **10** | (A;A) | non-inflamed | duodenum | M | 37 | 5B, C, E, F | |
| **11** | (A;A) | non-inflammed | terminal ileum | M | 23 | 5B, C, E, F | [5] |

**Table S2: characteristic of human serum samples**

| Patient ID | SARS-CoV-2 | BSI micro-organism |
|---|---|---|
| A | - | *E. faecium* |
| B | - | *P. aeruginosa* |
| C | - | *C. glabrata* |
| D | - | *C. glabrata* |
| E | - | *C. glabrata* |
| F | - | *E. faecalis* |
| G | - | *E. coli* |
| H | - | *E. coli* |
| I | - | *E. cloacae* |
| J | - | *S. aureus* |
| K | + | *E. faecium* |
| L | + | *C. dulinesis, S. aureus* |
| M | + | *S. aureus* |
| N | + | *P. aeruginosa* |
| O | + | *E. cloacae* |
| P | + | *E. faecalis* |
| Q | + | *E.coli* |
| R | + | *Serratia* |

**Table S3: Primers for RT-PCR**

| Gene | Forward Primer | Reverse Primer |
|---|---|---|
| Gapdh | TGGCCTTCCGTGTTCCTAC | GAGTTGCTGTTGAAGTCGCA |
| 18S | GTAACCCGTTGAACCCCATT | CCATCCAATCGGTAGTAGCG |
| Mx2 | CCAGTTCCTCTCAGTCCCAAGATT | TACTGGATGATCAAGGGAACGTGG |
| OasL2 | GGATGCCTGGGAGAGAATCG | TCGCCTGCTCTTCGAAACTG |
| Isg15 | GGTGTCCGTGACTAACTCCAT | TGGAAAGGGTAAGACCGTCCT |
| Zbp1 | AAGAGTCCCCTGCGATTATTT | TCTGGATGGCGTTTGATTGG |
| Mlkl | AATTGTACTCTGGGAAATTGCCA | TCTCCAAGATTCCGTCCACAG |
| Tnf | AGCCAGGAGGGAGAACAGAAAC | CCAGTGAGTGAAAGGGACAGAACC |
| Casp3 | ATGGAGAACAACAAAACCTCAGT | TTGCTCCCATGTATGGTCTTTAC |
| Casp8 | TGCTTGGACTACATCCCACAC | TGCAGTCTAGGAAGTTGACCA |
| Ripk3 | TCTGTCAAGTTATGGCCTACTGG | GGAACACGACTCCGAACCC |
